# DNA methylation responses to stress across different plant species

**DOI:** 10.1101/2023.05.10.540154

**Authors:** Morgane Van Antro, Slavica Ivanovic, Maarten Postuma, Lauren M. McIntyre, Wim H. van der Putten, Philippine Vergeer, Koen J.F. Verhoeven

## Abstract

- Changes in environments trigger plant DNA methylation responses, potentially modulating stress responses. Studies on stress-induced DNA methylation typically focus on single species, limiting our understanding of what are general and specific responses between species.
- Using reduced-representation bisulfite sequencing epiGBS, we compared DNA methylation stress-responses across seven plant species. Because methylation can be targeted at transposable elements (TEs) and because environmental plasticity may be particularly relevant in asexual species, we hypothesize that genome size and reproduction mode explain differences in methylation responses between species.
- We show that enrichment of environmentally-induced methylation responses in genes and TEs is a general feature across plant species. While previous studies have emphasized methylation responses in CHH-cytosines, we observed that cytosines in all sequence contexts (CG, CHG, CHH) are equally likely to respond to stress. Larger-genome species showed a higher proportion of stress-responding cytosines, and asexual species showed more cytosines with a strong methylation response to stress than sexually responding species.
- Our study highlights the role of TEs in methylome plasticity and identifies causes of variation in methylome plasticity. This provides guidance to extrapolating results from models to other plant species, and may contribute to better understanding of functionality of the response.

## Introduction

Plants are sessile organisms that constantly need to adapt to changes in their environment. As a result, plants have developed a wide range of mechanisms that enable them to cope in stressful conditions. Many of these mechanisms operate at the molecular level by modifying gene expression, which in turn induces changes in physiological and morphological traits (Sommer, 2020; Zhang *et al*., 2021b). Epigenetic modifications, such as DNA methylation, have been shown to regulate gene expression during normal plant development and functioning and are also thought to mediate plant stress responses by regulating gene expression induced during changes in environmental conditions (Herrera & Bazaga, 2013; Duncan *et al*., 2014; Zhang *et al*., 2018; vanden Broeck *et al*., 2018). In plants DNA methylation, defined as the addition of a methyl group to a nucleotide, occurs almost exclusively on cytosines, in three nucleotide contexts known as the CG, CHG and CHH cytosine context (where H = A, T or C). Each cytosine context is regulated by distinct but interconnected enzymatic pathways (Richards, 2006; Zhang *et al*., 2018; Gallego-Bartolomé, 2020). In addition to its role in modulating gene expression, the methylation of all three cytosine contexts within transposable elements (TEs) plays a prominent role in mediating TE activity (Zhang *et al*., 2018; Kumar & Mohapatra, 2021).

Although the functional role of environment-induced DNA methylation is not always evident (Secco *et al*., 2015; Bewick & Schmitz, 2017), it is often proposed that DNA methylation can translate environmental signals to modified gene expression profiles, thus acting as a regulating mechanism for the expression of phenotypic plasticity (Herrel *et al*., 2020; Skinner & Nilsson, 2021). Several studies have reported that cytosine contexts and genomic features are differentially impacted by stress and could thus play different roles in mediating stress responses (Zhang *et al*., 2018; Gallego-Bartolomé, 2020; Kumar & Mohapatra, 2021). Furthermore, while plant methylome responses to both biotic and abiotic stresses may be common, further studies suggest differences in how DNA methylation responds to stress both within and across plant species (Dubin *et al*., 2015; Galanti *et al*., 2022; Peña-Ponton *et al*., 2022). Conversely, methylation changes in CG context, and to a lesser extent CHG, are extremely stable and more often reflect genomic origin of an individual (Peña-Ponton *et al*., 2022; Ibañez *et al*., 2023). Opposing results have been reported in other plants species however ((López *et al*., 2022; Van Antro *et al*., 2022). What remains uncertain is to what extent such observed differences between species occur due to biological differences between study species or simply due to differences in experimental designs between studies. Thus, better insights in the generalities and specificities in DNA methylation stress responses across plant species are needed. A better understanding of these aspects of the methylation response to stress may help to establish how important environmentally-induced DNA methylation variations are in regulating stress responses.

Whole Genome Bisulfite Sequencing (WGBS) or Reduced Representation Bisulfite Sequencing (RRBS), are the golden standard techniques used to obtain high-resolution DNA methylation information in the genome (Laine *et al*., 2022). However, these techniques require the availability of a reference genome which are rarely available in plant species (Kress *et al*., 2022). Reduced-representation methylation screening techniques such as epiGBS (van Gurp *et al*., 2016; Gawehns *et al*., 2022) have been developed to circumvent the necessity of a reference genome for detecting DNA methylation variations. Thus, such techniques allow more flexibility in choosing plant species to compare. Such liberty in choosing plant species, allows for experiments to be designed to specifically examine biological characteristics such as reproduction mode and genome size in order to determine whether these factors influence epigenetic stress responses.

Lacking the same potential for genetic adaptation that sexual species have, clonal species have been predicted to depend more strongly on phenotypic plasticity than sexually reproducing organisms to cope with spatial and temporal environmental heterogeneity (Lynch, 1984; Massicotte & Angers, 2012). Thus, it has been hypothesised that environmental plasticity in epigenetic mechanisms is particularly well-developed in asexual taxa (Castonguay & Angers, 2012; Verhoeven & Preite, 2014; Zhang *et al*., 2021a; Sammarco *et al*., 2022; Wibowo *et al*., 2022). Genome size has been shown to guide the natural methylation landscape of plant species, mainly via variation in abundance of transposable elements (Seymour *et al*., 2014; Alonso *et al*., 2015; Niederhuth *et al*., 2016). TEs are methylated in all three cytosine contexts and respond to environmental stresses (Casacuberta & González, 2013; Miousse *et al*., 2015; Negi *et al*., 2016; Hämälä *et al*., 2022), leading to the hypothesis that larger genomes express stronger methylome plasticity. A relation between genome size and methylome responses to environmental stress has, however, not yet been demonstrated.

In the present study, we describe and compare the DNA methylation response to stress between plant species that differ in genome size and in reproduction mode, with the aim to gain more insight in both the generalities and species-specific aspects of plant methylome responses to stress. Using epiGBS, we screen for DNA methylation patterns in seven different plant species after exposure to drought and/or salicylic acid treatment. Plant methylomes have been shown to respond to both drought and salicylic acid in previous studies (Arora *et al*., 2022; J. Liu & He, 2020b; M. Sun *et al*., 2022). We ask the following questions: 1) what aspects of the DNA methylation response to stress are conserved between plant species (reflecting a general response)? 2) What aspects of the response show clear differences between species (reflecting species-specific responses)? 3) To what extent can observed differences in responses be explained by genome size or reproduction mode differences between species?

## Material and Methods

### Species Choice

Eight diploid plant species were chosen based on their reproduction mode and genome size. Species were either inbred selfing sexuals or could reproduce clonally (Table 1). The genome size of the species ranged between 158 and 3234 Mbp (Table 1). Sexually reproducing species were *Arabidopsis thaliana, Capsella rubella, Mimulus laciniatus* and *Bromus tectorum*. Clonally reproducing plants were *Fragaria vesca, Rorripa austriaca, Linaria vulgaris* and *Solidago canadensis*. While the chosen clonal species are able to reproduce sexually, clonal reproduction via rhizomes or stolons is common for all of these species. Due to poor DNA quality and low sequencing read output of multiple samples, *Fragria vesca* was dropped from the analysis. By selecting inbred selfing sexual species, we attempted to minimise effects of segregating genetic variation between replicate plants, allowing for a fair comparison of DNA methylation variation with the clonal species.

**Table 1:**
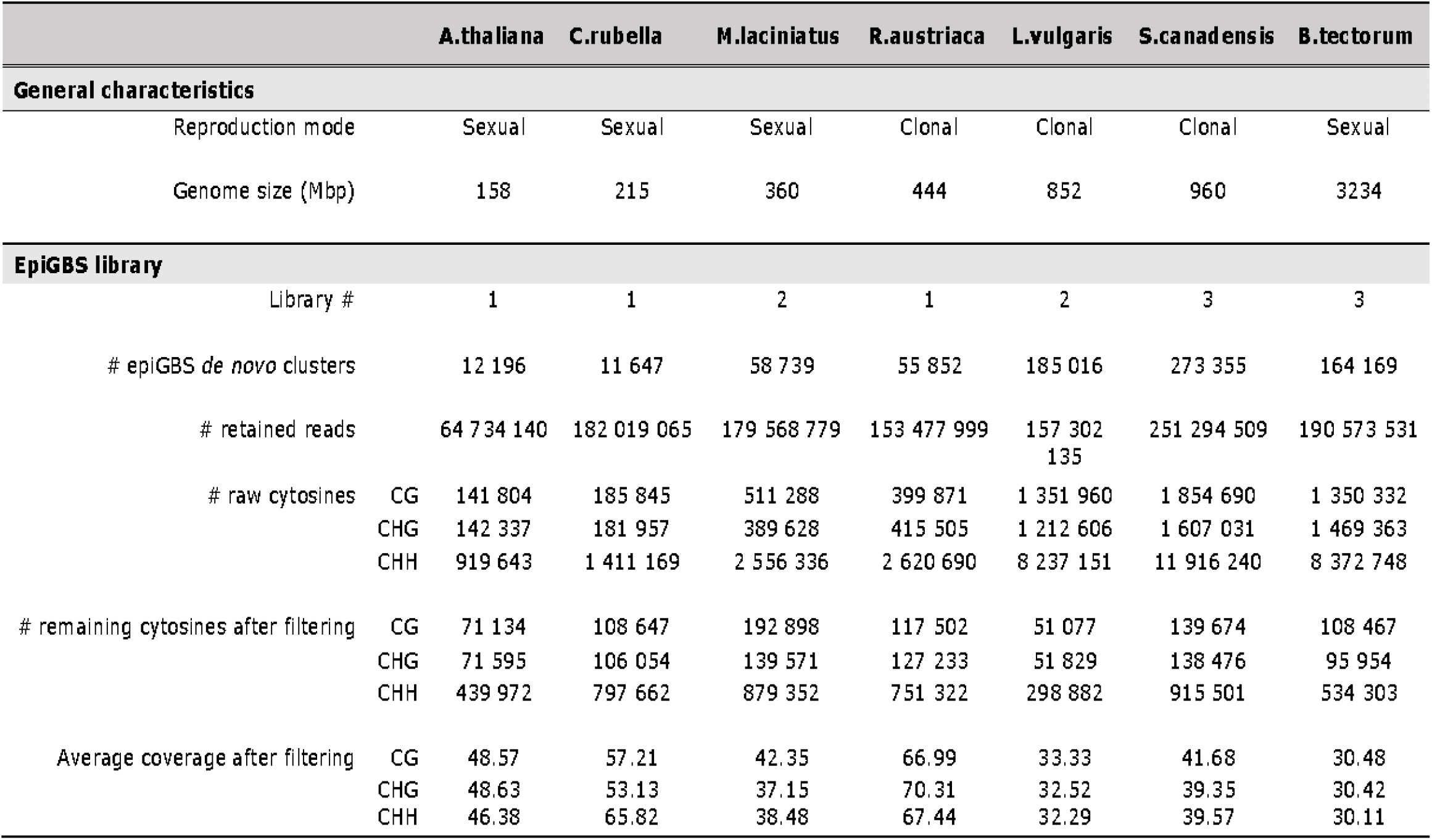
Information for each species on reproduction mode, genome size, within which epiGBS library it was sequenced, the number of de novo clusters generated, the number of raw cytosines detected (CG, CHG and CHH cytosine context), the of cytosines left after filtering (CG, CHG and CHH cytosine context) and the average coverage after filtering (CG, CHG and CHH cytosine context).

### Plant Material and Germination

Seeds (see Supplementary S1) were stored, until gemination, in darkness at 15°C and 30% relative humidity. All species were germinated following the same protocol: Around 30 seeds were sterilised by placing them in 8% bleach for 5 minutes and then rinsed with demineralized water three times. Seeds were then stratified in the dark at 4°C for 10 days by placing them on moist, sterilised filter paper. After stratification, seeds were placed on new, moist, sterilised filter paper, in Petri-dishes and allowed to germinate in climate cabinets (16h/8h light/dark; 21°C/16°C). Once the cotyledons fully emerged, seedlings were transferred to plastic containers filled with seedling soil (*ZST-D*, Zaai/Stek basis) and placed in climate cabinets (16h/8h light/dark; 21°C/16°C). When 80% of individuals had reached the 3rd leaf stage, seedlings were vernalized at 4°C in the dark for 7 days. Seedlings were then placed back into the climate cabinet for 1 day before planting 20 3-4 leafed seedlings in individual pots filled with 3:1 potting soil:pumice mixture (*BPO-D*, Lensli substrate), adding 0.5ml of Osmocote slow-release fertiliser granules (ICL, Osmocote Extract Mini 5-6M). The top soil was covered with coarse sand to protect from pest contamination and avoid excessive growth of mosses. Pots were then transferred to a greenhouse with the following conditions: Set point day: 06:00-22:00, 21°C (−1/+2-4°C); Set point night: 22:00 - 06:00, 16°C (−1/+2°C). High pressure Sodium lamps (Son-T, 600W Philips GP) were used when insufficient daylight was present, plants thus received a minimum of ∼225 u.moll/m2/s PAR. Average level of humidity inside the greenhouse was 60% Rh (−5/+5% Rh).

### Experimental Design

For each species, individuals were grown in controlled greenhouse conditions as described above until 80% of individuals had reached the 5-6 leaf stage. Individuals of similar developmental stages were then selected and randomly assigned to one of four treatment group: Controls (CTL); Drought (D); Salicylic Acid (SA) or a combination of Drought + Salicylic Acid (DSA). Each group consisted of 4-5 replicates, leading to an experimental design of maximum 20 individuals per species. Individual plants were fully randomised within a species. Species were exposed to episodic stress cycles during their vegetative growth phase (Fig.1). One stress cycle consisted of a two-week stress period followed by a one-week recovery period. The stress cycles were repeated until 80% of the plants within one species showed signs of bolting (sexual species) or after three stress cycles (clonal species). Because the time of bolting differed between sexually reproducing species, the number of stress cycles ranges from 1-6 in these species (average ∼3 cycles, which equals the number of cycles applied to all clonal species).

**Figure 1:**
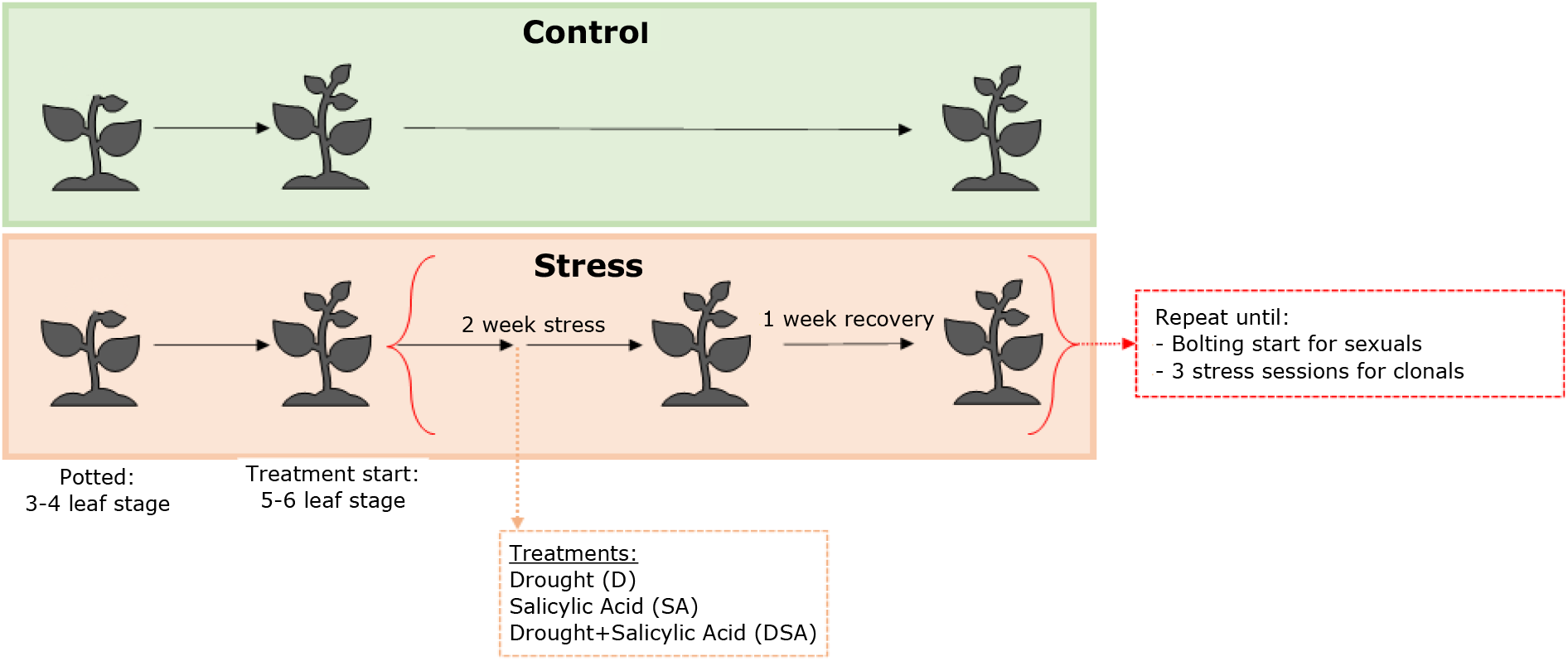
Overview of experimental design. 7 plant species were exposed to either drought, salicylic acid or a combination of drought and salicylic acid. Treatments were applied over stress cycles which consisted of applying the stress for a period of 2 weeks before allowing for a one week recovery. Stress cycles were repeated until species that reproduce sexually showed signs of bolting or for three stress cycles for clonal species.

The environmental treatments were applied as follows: Control individuals were maintained under greenhouse conditions as described above and were watered to full soil capacity three times a week. Drought stress was applied by withholding water for a period of two weeks, immediately followed by a one-week recovery period during which individuals were watered as per the control group. Salicylic acid was applied once a week during the two-week stress periods, by pipetting 1mM of salicylic acid solution (Sigma-Aldrich) on 20% of the leaves and spreading the solution across the top and bottom surfaces of the leaves. Depending on leaf size of the species, between 20-100 μl of 1% salicylic acid was applied per leaf. Watering regime and greenhouse conditions were as per the control group.

Because of differences between species in germination timing and growth, the experiment was not executed as one single, fully randomised experiment. Instead, each species was grown separately as an individual experiment, where all species experiments were started within an 8-week period. As a result, all lab work and downstream bioinformatic and statistical analyses were performed separately per species.

### epiGBS Library Construction

#### Sampling and DNA extraction

After the last stress cycle during the vegetative growth, a one-week recovery period was allowed before tissue sampling. Sampling for DNA methylation analysis consisted of collecting 6-8 circular leaf punches (Ø 8 mm). Due to narrow leaf shape, *B.tectorum* and *L. vulgaris* were sampled differently, with the top half of the second newest leaf being sampled for *B.tectorum* and 2-3 leaves from the tallest inflorescence branch sampled for *L. vulgaris*. Leaf samples were flash-frozen in liquid nitrogen and stored in -80C until further analysis. To prepare samples for DNA extraction, the frozen leaf material was homogenised using a Qiagen TissueLyser II with the use of two stainless steel beads (2 × 45 seconds at a frequency of 30/s). DNA isolation was then performed using the Macherey-Nagel NucleoSpin Plant II kit (using the lysis buffer PL1 for all species) ending with samples being diluted to 30 ng/ul of DNA.

#### epiGBS library preparation

Eight epiGBS libraries (van Gurp *et al*., 2016) were prepared (one for each species) following the epiGBS2 protocol (Gawehns *et al*., 2022). In brief: Methylation insensitive restriction enzymes AseI and NsiI were used to digest individual DNA samples. The digested DNA was then ligated with hemi-methylated barcoded adapter pairs (each barcode pair containing 4-6 sample specific nucleotides, followed by a Unique Molecular Identifier of three nucleotides and an unmethylated cytosine) allowing for all samples of one species to be multiplexed together. Multiplexed samples were then concentrated before the removal of smaller fragments (<60bp) using the NucleoSpin Gel & PCR cleanup Kit. DNA fragments sized between 300-400bp were then selected using 0.8X SPRIselect magnetic beads. Samples were bisulfite converted using EZ DNA Methylation-Lightning kit. The converted DNA was then PCR-amplified, concentrated, cleaned and size-selected once more. Finally, in order to ensure a balanced distribution of reads per species, taking into account differences in genome size and final DNA concentrations, 2-3 species were equimolarily pooled together into a final epiGBS library (Library 1: *A.thaliana, C.rubella, L.vulgaris*; Library 2: *M.laciniatus, F.vesca, R.austriaca*; Library 3: *S.canadensis, B.tectorum*). The final epiGBS libraries were sequenced paired-end (PE 2×150bp) with a 12% phiX spike on one sequencing lane per library using an Illumina HiSeq X sequencer.

#### epiGBS analysis

While some species have a published referenced genome, all species were analysed using the ‘*de novo*’ branch of the epiGBS2 pipeline as to avoid species-specific analysis biases. All sequencing data was processed per species through the epiGBS2 pipeline (Github commit 125eebe) (Gawehns *et al*., 2022), using the *de novo* branch’s default parameters (Sequencing identity of 0.97 for the last clustering step; Minimum cluster depth of 10; Maximum cluster depth of 10000).

To ensure that the libraries were of high quality, we performed a quality control of the sequencing data of each species. We looked at the total number of cytosines obtained before and after demultiplexing and the number of epiGBS *de novo* clusters constructed (Table 1). One species, *F. vesca* showed a very low number of reads and so was removed from the analysis. Subsequently, the following filtering steps were applied to the raw demultiplexed sequencing data: Firstly, any sample for which the number of reads was below 10% of the average total number of reads for that species was removed from the analysis (one sample in *C.rubella* and one sample in *R.austriaca*). Next, a minimum 10X coverage threshold was applied, and the 0.1% sites with the highest coverage were excluded from the analysis. The coverage distribution before and after filtering, for each species, can be found in Supplementary S2. Subsequently, only cytosines present in at least 80% of all samples (irrespective of stress group) were considered in the analysis (See Table 1 for values).

### Annotation

The *de novo* epiGBS reference sequences of each species were individually annotated by homology (Supplementary S3) in the following manner: In order to obtain genomic information on TEs and repeat regions, the epiGBS reference fragments were annotated through REPEATMASKER (Embryophyta as the reference species collection, version 4.1.2, Dfam database 3.6 uncurated) (Smit, Hubley, Green, https://repeatmasker.org). Homologous gene information was obtained by using the DIAMOND database (version 2.0.14) (Buchfink *et al*., 2014). Both programs were run using default settings.

### Global DNA methylation level

Global methylation level of each species and cytosine context was determined in individuals from the control group and was calculated as the average per-cytosine methylation level for each individual sample. Knowing that normality (Shapiro-Wilk’s test) and homogeneous variance (Levene’s test) assumptions were met; effects of drought, salicylic acid, and the DSA interaction on global methylation level were tested for each cytosine context using a two-way ANOVA model (Average methylation levels ∼ Drought main effect * Salicylic main effect; *aov* function, base R v3.6.2). In order to detect if a significant correlation exists between genome size and average methylation levels, a Pearson correlation test (*cor* function, base R v3.6.2) was applied for each cytosine context, using the average methylation level of each species as a data point.

### Differentially Methylated Cytosines (DMCs)

In order to identify differentially methylated cytosines (DMCs), a beta-binomial regression model was fitted to each individual cytosine using the R package DSS (Park & Wu, 2016), to test for effects of drought, salicylic acid, and their interaction, on cytosine methylation level. The model used an arcsine link function, Wald testing and no smoothing was applied. Cytosines with a mean methylation level across all samples of <3% or >97% were removed from the analysis, thus removing the majority of cytosines that showed little variation between samples. A false discovery rate (FDR) threshold of 0.05 was applied to account for multiple testing.

When comparing the proportion of cytosines that are differentially methylated between species and cytosine contexts, we refrained from FDR correction as correcting for FDR would create a genome size-related bias (the multiple testing penalty is larger for cytosines in large genomes and for CHH cytosines (for which there are relatively more compared to CG and CHG cytosines, resulting in reduced power to detect DMCs). Instead, we labelled cytosines as ‘responsive cytosines’ when their raw p-value was <0.05 and the methylation difference between experimental groups was >5 percentage points difference. We then compared the proportion of ‘responding cytosines’ (among all cytosines tested) between species and between cytosine contexts. A logistic regression test was used to test for a significant relationship between genome size (log transformed) and the proportion of cytosines that respond to stress (proportion of responding cytosine ∼ log(Genome size)+ Stress; g*lm* function, quasi-binomial famlily, base R v3.6.2). For this model, the species-level estimates for each context of the proportion of responding cytosines were used as independent data points.

We labelled the subset of DMCs that showed a minimum 20 percentage point methylation difference between experimental groups as ‘strongly responding’ cytosines. In order to test if reproduction mode significantly influenced the species-level proportion of strongly responding cytosines, a logistic regression test was performed, using the species-level methylation estimates for each cytosine context and stress as independent data points (ratio of strongly responding DMCs ∼ Stress + Cytosine context + Reproduction mode + Reproduction mode/Species, *glm* function, quasi-binomial family; base R v3.6.2). Per-cytosine differences in DNA methylation between treatment groups were calculated as the difference in average per-cytosine methylation level between treatment groups.

To test if DMCs were enriched in a specific genomic feature (relative to random expectation based on the representation of each feature in the data set) an enrichment analysis was performed using Fisher Exact test (*phyper* function, base R v3.6.2). A genomic feature was considered significantly enriched in DMCs with p-value below 0.05. Fold change was calculated based on the following formula:

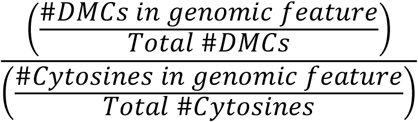

## Results

### ‘Static’ DNA methylation patterns across species

The ‘static’ DNA methylation landscape, as observed in control plants, varied greatly between species both in percentage and context of methylation, although concordant aspects were found. In all species, average global methylation levels were highest in CG context (18.48-77.42%), followed by CHG (4.81-39.8%) and then CHH (3.01-10.4%) (Fig.2B). A significant correlation between genome size and methylation levels was detected in the CG and CHG cytosine contexts (Fig. 2). CHH methylation levels were not significantly correlated to genome size but this was primarily due to the strikingly low methylation level in *B. tectorum* as the correlation became significant if *B. tectorum* was removed from the analysis (p value 0.0078). Additionally, for all species, cytosine methylation levels showed a bimodal distribution with most cytosines either having relatively high or low methylation levels (Fig. 2A). Species with smaller genomes showed lower average per-cytosine DNA methylation levels with 75-70% of CG or CHG cytosines having methylation levels less than 5%. This pattern inverted in species with larger genomes with the majority of CG cytosines being highly methylated (21-76% of CG or CHG cytosines having methylation levels higher than 80%). CHG methylation showed a similar trend as CG methylation but with a larger proportion of cytosines with intermediate methylation levels. Individual CHH cytosine methylation levels were relatively low in all species, with the majority of cytosines being unmethylated (close to 95% of cytosines in all species). Some deviations from this general genome size trend were however observed: *M. laciniatus* showed comparatively high cytosine methylation levels in all cytosine contexts, despite having a relatively small genome. Concurrently, *B. tectorum* showed relatively low methylation levels in CHH context, given its large genome size.

**Figure 2:**
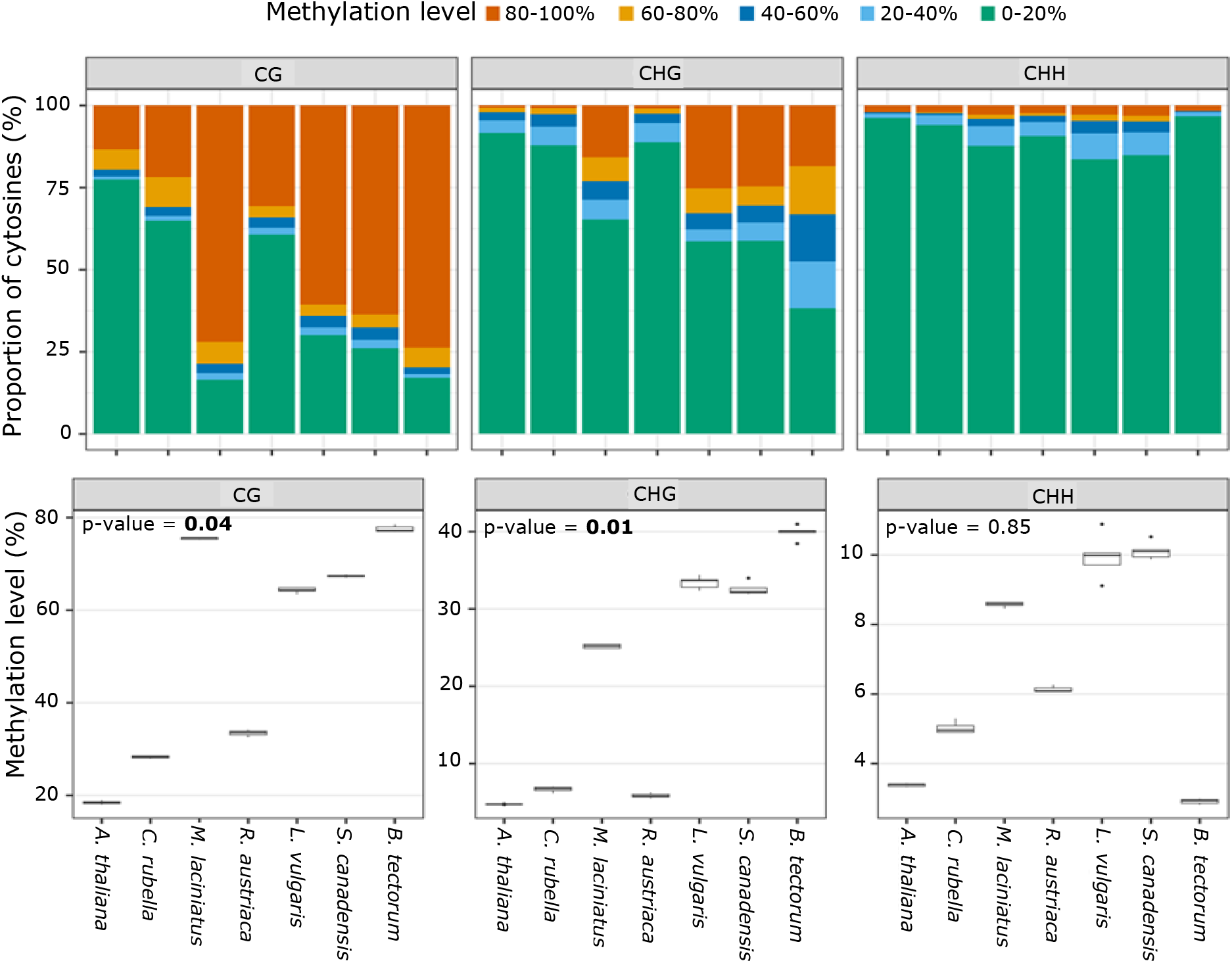
‘Static’ Methylome across plant species. A) Distribution of per-cytosine methylation levels, across species and per cytosine context (CG, CHG and CHH). DNA methylation levels were calculated as the mean value across all samples of per-cytosine methylation estimates. B) Average global methylation levels per plant species and per cytosine context (CG, CHG and CHH). A Pearson correlation test on the species-level average values was used to evaluate the association between genome size and methylation levels. Significant correlations (p value < 0.05) are shown in bold.

### Stress responses in DNA methylation

#### Genome-wide Methylation Levels

A significant effect of a stress on global methylation levels was detected in all targeted plant species for at least one stress and in at least one of the three cytosine contexts. This global stress response the DNA methylation levels in all plant species indicates that methylome stress-responses are typically not limited to a few cytosines but occur throughout the genome. Across species, responses due to drought were more common than responses to SA or due to an interaction between the two stresses (Fig.3, Table 2).

**Figure 3:**
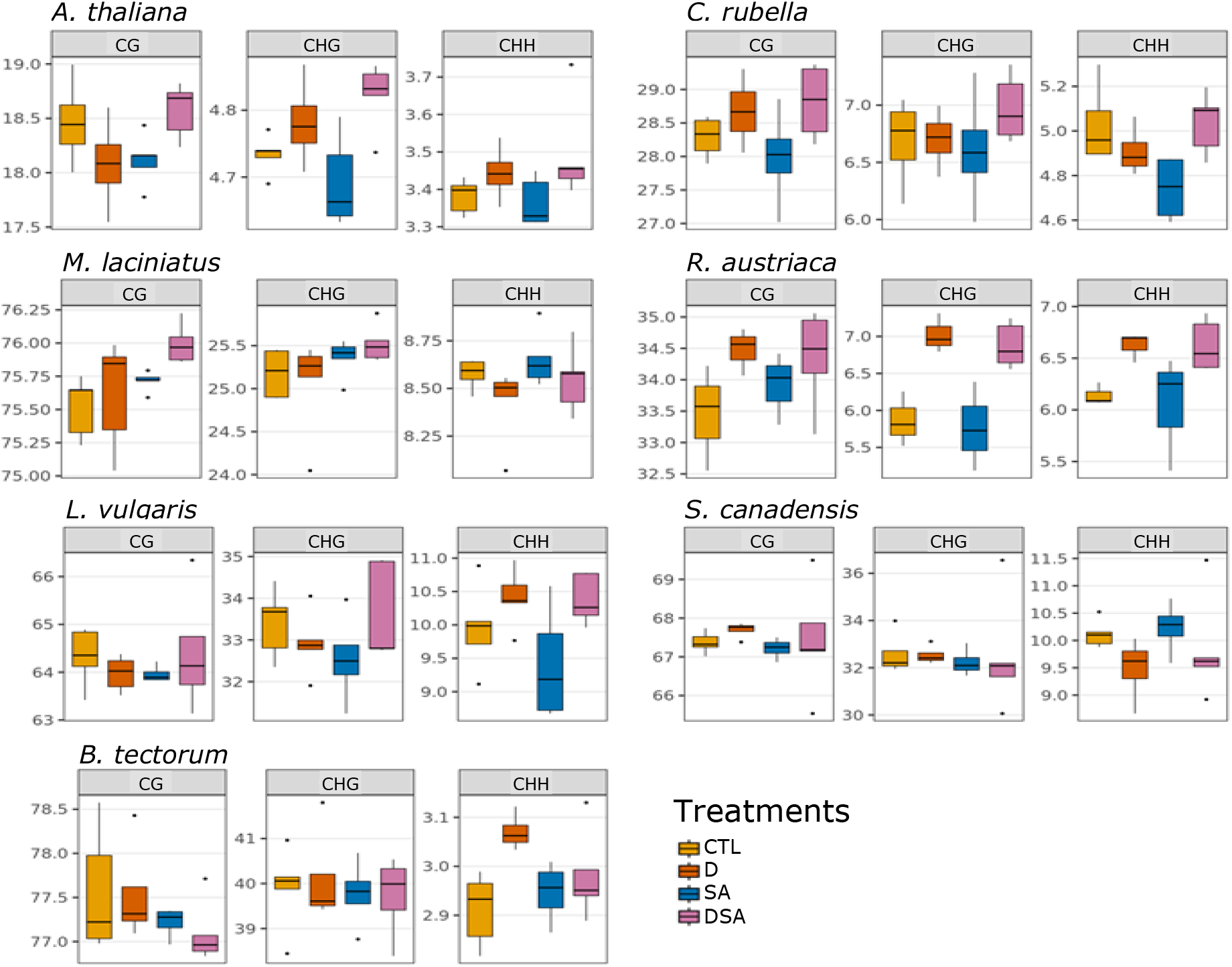
Average genome-wide DNA methylation levels across plant species per treatment group and per cytosine context, as affected by stress treatment. Plants species were grown in identical growth conditions and exposed to either Drought (D); Salicylic Acid (SA), a combination of both stresses (DSA) or no stress (CTL). Effects were considered significant when P-values < 0.05 (shown in bold).

**Table 4.2:**
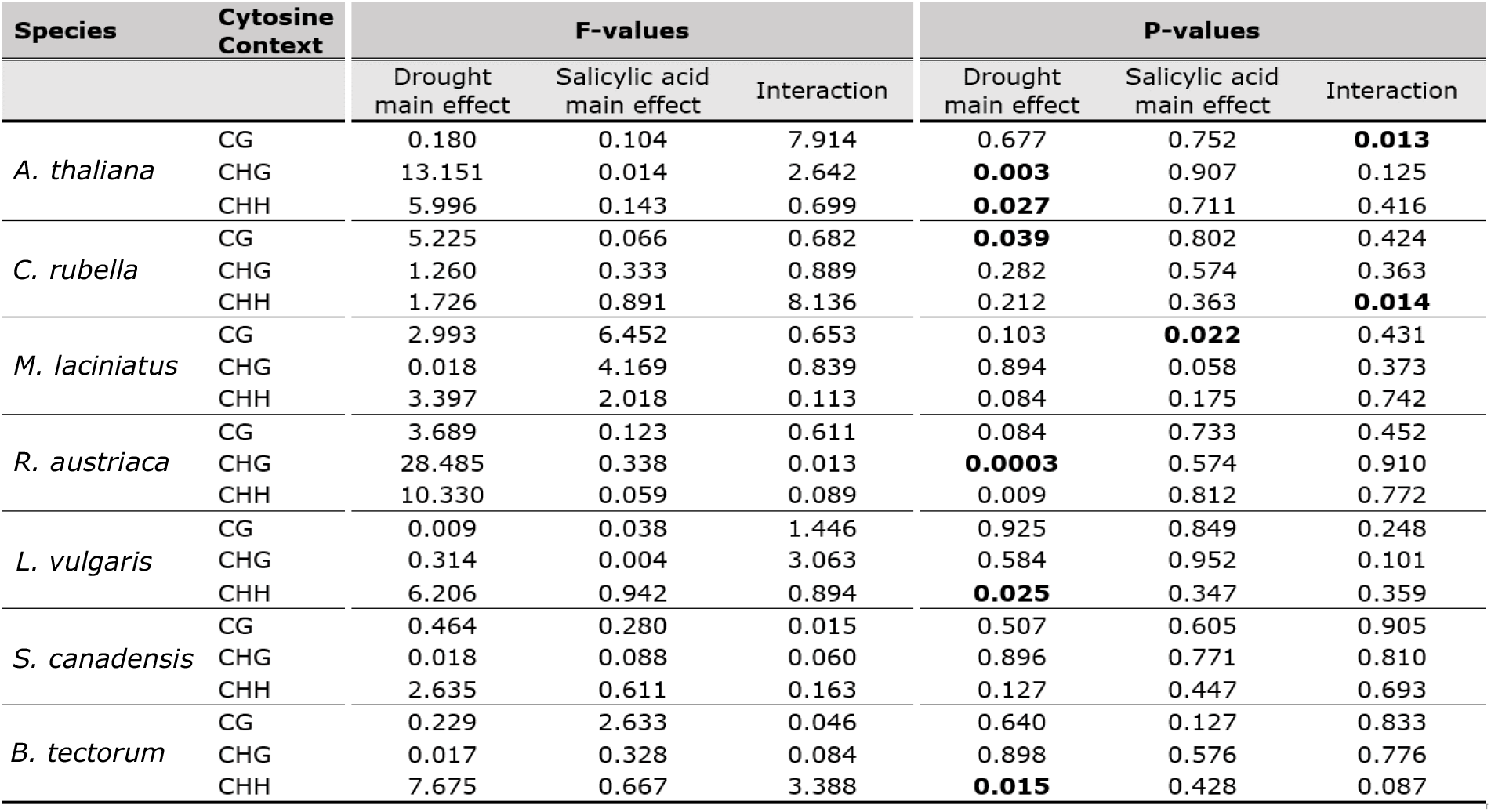
Two-way ANOVA table. Model tested the effect of drought main effect, salicylic acid main effect and the interaction on global methylation levels. Effects were considered significant when P-values < 0.05 (shown in bold).

| Species | Cytosine Context | F-values |  |  | P-values |  |  |
| --- | --- | --- | --- | --- | --- | --- | --- |
|  |  | Drought main effect | Salicylic acid main effect | Interaction | Drought main effect | Salicylic acid main effect | Interaction |
| *A. thaliana* | CG | 0.180 | 0.104 | 7.914 | 0.677 | 0.752 | **0.013** |
|  | CHG | 13.151 | 0.014 | 2.642 | **0.003** | 0.907 | 0.125 |
|  | CHH | 5.996 | 0.143 | 0.699 | **0.027** | 0.711 | 0.416 |
| *C. rubella* | CG | 5.225 | 0.066 | 0.682 | **0.039** | 0.802 | 0.424 |
|  | CHG | 1.260 | 0.333 | 0.889 | 0.282 | 0.574 | 0.363 |
|  | CHH | 1.726 | 0.891 | 8.136 | 0.212 | 0.363 | **0.014** |
| *M. laciniatus* | CG | 2.993 | 6.452 | 0.653 | 0.103 | **0.022** | 0.431 |
|  | CHG | 0.018 | 4.169 | 0.839 | 0.894 | 0.058 | 0.373 |
|  | CHH | 3.397 | 2.018 | 0.113 | 0.084 | 0.175 | 0.742 |
| *R. austriaca* | CG | 3.689 | 0.123 | 0.611 | 0.084 | 0.733 | 0.452 |
|  | CHG | 28.485 | 0.338 | 0.013 | **0.0003** | 0.574 | 0.910 |
|  | CHH | 10.330 | 0.059 | 0.089 | 0.009 | 0.812 | 0.772 |
| *L. vulgaris* | CG | 0.009 | 0.038 | 1.446 | 0.925 | 0.849 | 0.248 |
|  | CHG | 0.314 | 0.004 | 3.063 | 0.584 | 0.952 | 0.101 |
|  | CHH | 6.206 | 0.942 | 0.894 | **0.025** | 0.347 | 0.359 |
| *S. canadensis* | CG | 0.464 | 0.280 | 0.015 | 0.507 | 0.605 | 0.905 |
|  | CHG | 0.018 | 0.088 | 0.060 | 0.896 | 0.771 | 0.810 |
|  | CHH | 2.635 | 0.611 | 0.163 | 0.127 | 0.447 | 0.693 |
| *B. tectorum* | CG | 0.229 | 2.633 | 0.046 | 0.640 | 0.127 | 0.833 |
|  | CHG | 0.017 | 0.328 | 0.084 | 0.898 | 0.576 | 0.776 |
|  | CHH | 7.675 | 0.667 | 3.388 | **0.015** | 0.428 | 0.087 |

#### Proportion of stress-responsive cytosines

Interestingly, within a species, cytosines in all three sequence contexts were equally likely to show a methylation response to the stress treatments (Fig. 4). The estimated proportion of responding cytosines varied between 5.35-11.70% for cytosines in CG context (average of 8% across species), 5.42-15.26% for cytosines in CHG context (average of 8.64%) and 4.14-12.59% for cytosines in CHH context (average of 8.13%). We found a significant relationship between genome size and the proportion of cytosines that respond to a stress in the CHH and CHG cytosine context (p-value = 0.047 and 0.039 for CHG and CHH respectively) but not for CG (p-value = 0.12). These results have to be interpreted with some caution: responding cytosines were defined as showing significant differential methylation of p-value < 0.05 and having a 5-percentage point difference. No multiple testing correction was applied, as such correction would create different significance thresholds for different species and sequence context, making a comparison among species difficult (see Material and Methods). Consequently, the estimated values include a fraction of false positives (maximum expectation 5% of all tested cytosines, but presumably less because significant cytosines also needed to show at least a 5-percentage point difference in methylation levels to be labelled as a ‘responding cytosine’). Because the proportion of false positives is expected to be the same for all species and contexts, we consider that the absolute level of the estimates is inflated by false positives, but the differences in the estimates between species and contexts can be interpreted. Two additional results are noteworthy: First, the proportion of responding cytosines in *A. thaliana* was low compared to other species, irrespective of stress and cytosine context (5.35-5.61% for CG; 5.42% for CHG and 4.14-4.48% for CHH). Second, a larger fraction of cytosines responded to the stress in some species, in particular *R. austriaca* in response to drought stress and *L. vulgaris* in response to SA.

**Figure 4:**
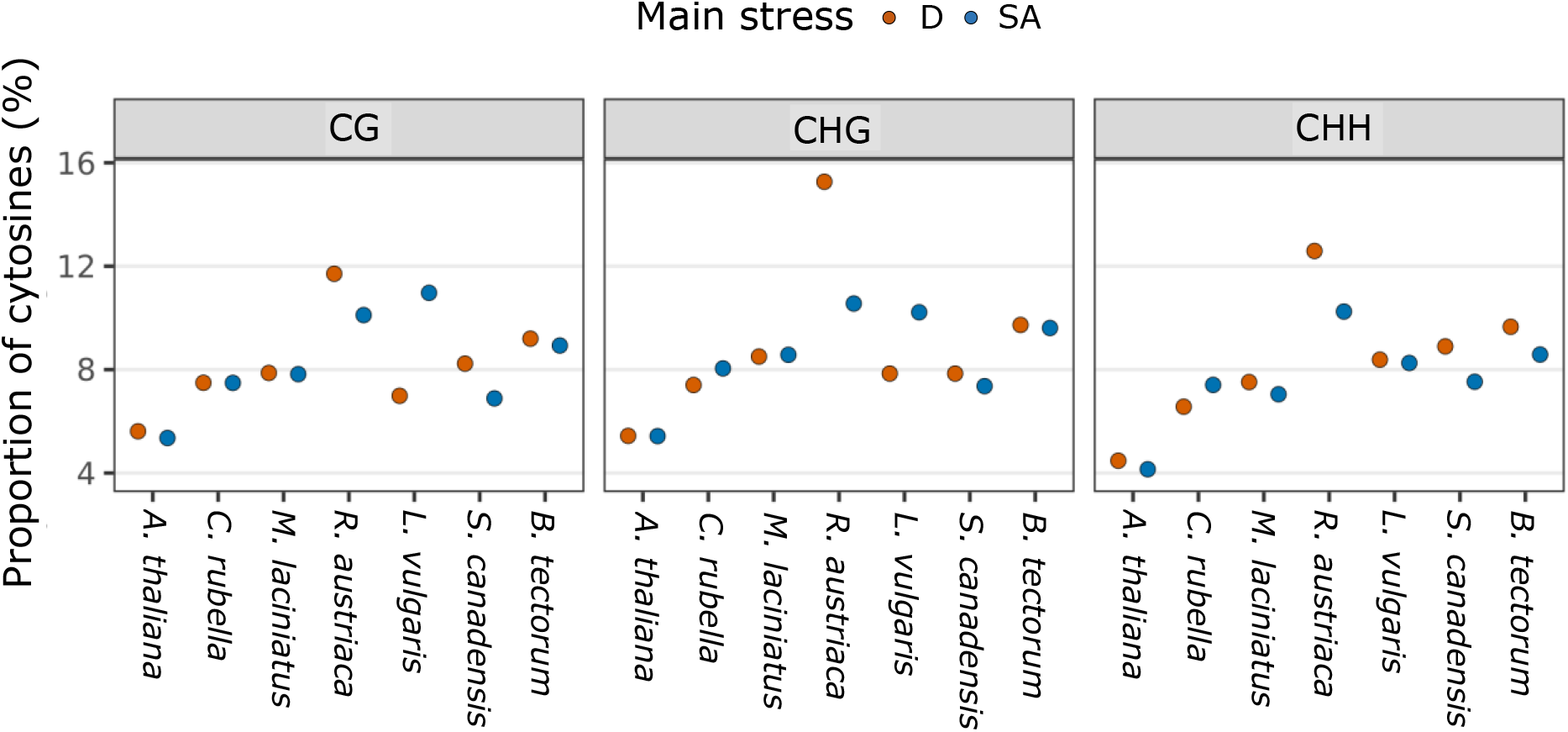
Proportion of all tested cytosines that show a response to stress, calculated for each species per stress and per cytosine context. Cytosines were considered responding to a stress treatment if the percentage point difference in methylation levels was >5% and the P-value for the test of the main stress effect on methylation level was < 0.05. Species are ordered in increasing order of genome size.

#### Differentially Methylated Cytosines (DMCs)

##### Generic vs stress-specific DMCs

The methylation response to stress can show high specificity (different stresses trigger different methylation responses) or can be generic (different stresses trigger the same methylation responses). In Fig.5, we distinguish between DMCs that show only a significant effect of drought, DMCs that show only significant effect of SA, and DMCs that show an effect of both treatments and/or that show a significant interaction effect between the two treatments (meaning that the methylation response to one stress is conditional on the presence of another stress). We interpret the first two as stress-specific and the latter as reflecting a generic response. For all species and cytosine contexts, the methylome response to the drought and SA treatments experimental treatments consisted of both a treatment-specific component and a more generic stress response (Fig.5). Overall, similar proportion of stress-specific DMCs were induced in each plant species, with the exception of some species where clear differences were detected (Fig.5). In most species and over all cytosine contexts, the proportion of generic DMCs ranged between 21-61% for CG DMCs, 23-61% for CHG DMCs and 29-61% for CHH DMCs. Yet, all species induced stress-specific DMCs, with some species showing a considerably stronger response to a particular stress (*L. vulgaris* and *A. thaliana* in response to SA; *R. austriaca* and *S. canadensis* in response to drought.

**Figure 5:**
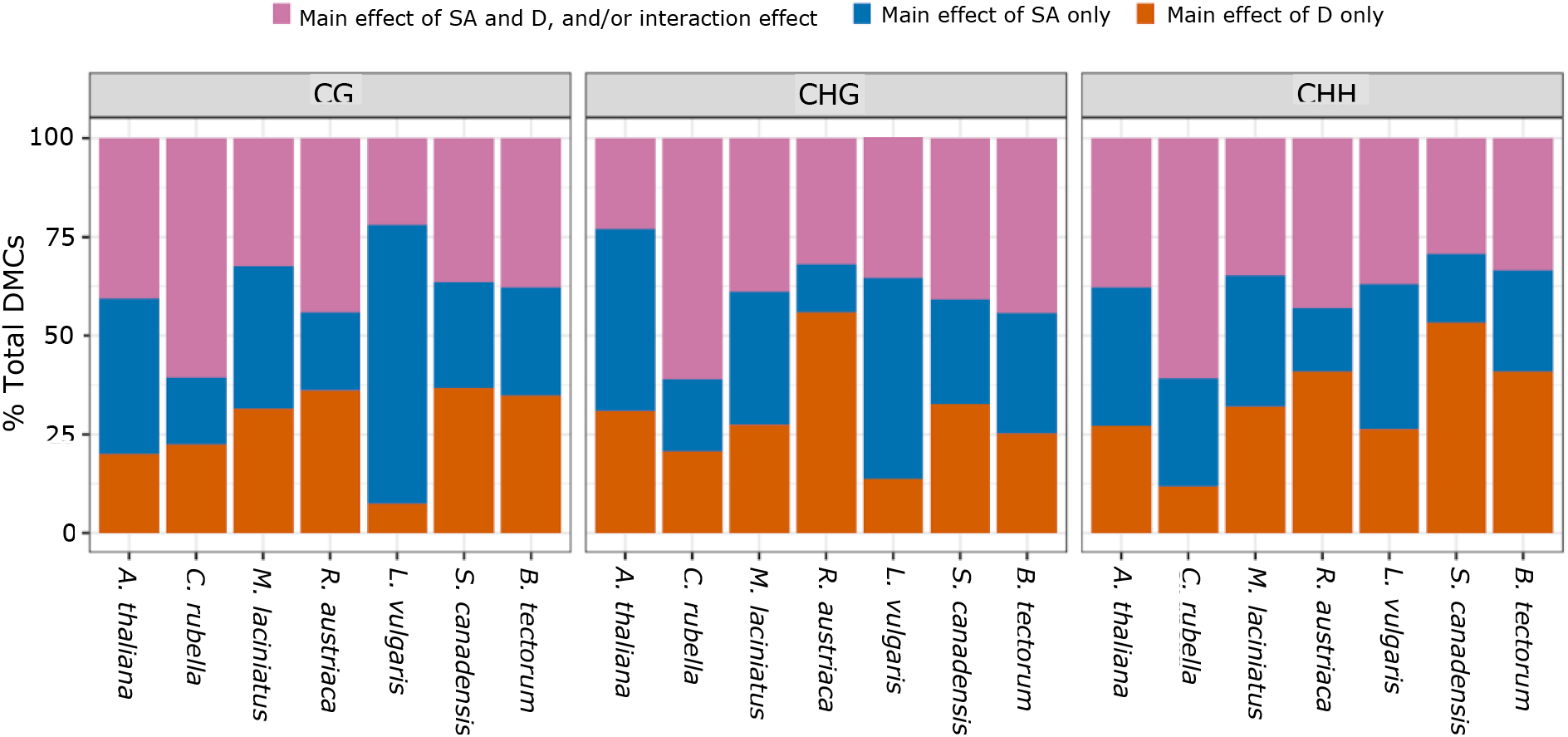
Proportion of DMCs that are responsive to either one of the stresses or to both stresses, per species and per cytosine context. DMCs are classified as having only a significant main effect of drought (orange bar), only a significant effect of SA (blue bar), or a significant effect of both stresses and/or a significant interaction effect between the stresses (purple bar). Cytosines were labelled as DMCs at an FDR threshold of 0.05.

##### Strongly stress-responding DMCs

While the majority of significant DMCs showed only a modest change in DNA methylation level, across all species between 0-42.5% of CG DMCs, 0-36.7% of CHG DMCs and 0-25.2% of CHH DMCs were classified as strongly responding to a stress (showing a methylation percentage point difference of 20% or higher between stress and control groups; Fig.6). An extreme case was *A. thaliana*, with less than 1% of identified DMCs showing strong methylation differences when stressed. Using species-level estimates as independent data points, the proportion of strongly responding cytosines was significantly different between sequence contexts (p-value <0.001, with CHH showing a smaller proportion) but not between stresses (p-value >0.5). Interestingly, for both stresses, a significant effect of plant reproduction mode was detected: clonally reproducing species (*R. austriaca, S.canadensis* and *L.vulgaris*) induced a larger portion of strongly responding DMCs compared to the sexually reproducing species (p-value <0.005). This result was obtained either with or without including *A. thaliana* in the analysis.

**Figure 6:**
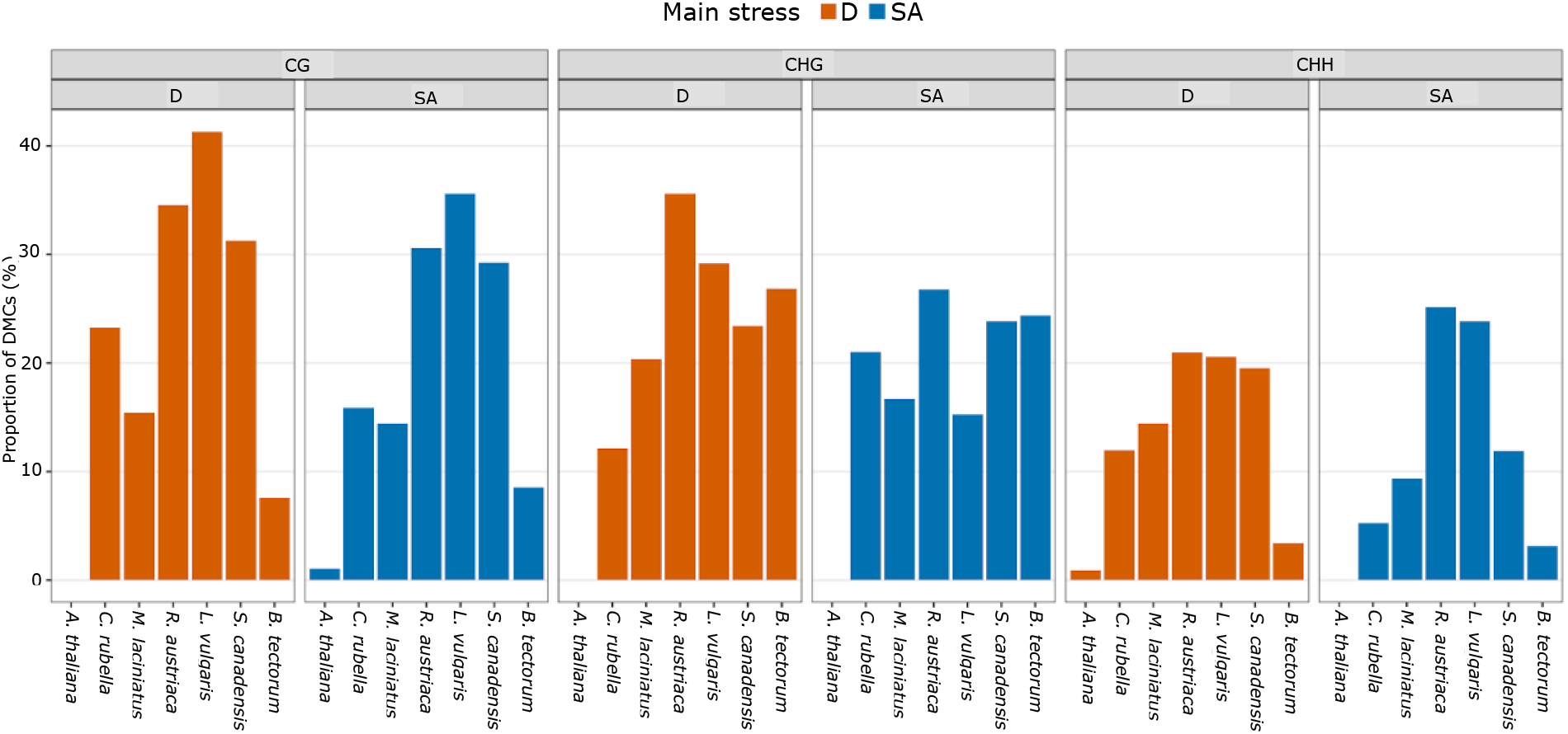
Proportion of DMCs that show a strong methylation response to a stress, calculated for each species and cytosine context. DMCs were considered strongly responding when the test of the main effect of the stress treatment was significant at an FDR < 0.05 threshold and the methylation difference was >20 percent points between the control and stress experimental groups.

##### Enrichment of DMCs within genomic regions

The majority of identified DMCs were detected within unannotated regions of the genome (between 38-86% of DMCs, depending on the species, stress and cytosine context; Supplementary S4). The remainder of the identified DMCs were found within genes (between 11-58 % of CG DMCs; 10-39% of CHG DMCs and 10-34% of CHH DMCs), with a relatively very small percentage of DMCs detected in TEs and repeat regions, for all species.

We observed significant enrichment of DMCs in genes and transposable elements (Fig.7). Specifically, in all species and for both stresses, CG DMCs showed a significant 1.1 to 2.2-fold enrichment in genes, with CHG and CHH DMCs being consistently enriched within TEs (by 1.2 to 3.1-fold for CHG and 1.3 to 3.2-fold for CHH). The only exception was *B. tectorum* which did not show an enrichment of CHH DMCs in TEs in response to SA. In addition to these general patterns, smaller-genome sized species (*A. thaliana, C. rubella* and *R. austriaca*) showed enrichment of CG-DMCs TEs. However, *M. laciniatus* once again behaved similarly to species with larger genomes (see point ‘DNA methylation patterns across plant species’), with no enrichment of CG DMCs in TEs detected, despite having a relatively small genome. Certain species showed distinct enrichment patterns that were not shared by other species, such as *S. canadensis* and *R. austriaca* showing strong enrichment of CHG-DMCs in genes in response to drought and SA, respectively, and *A. thaliana* showing significant enrichment of CHH-DMCs in genes. Repeat elements were also regularly enriched in DMCs, in particular in the CHH cytosine context and when species were exposed to drought.

**Figure 7:**
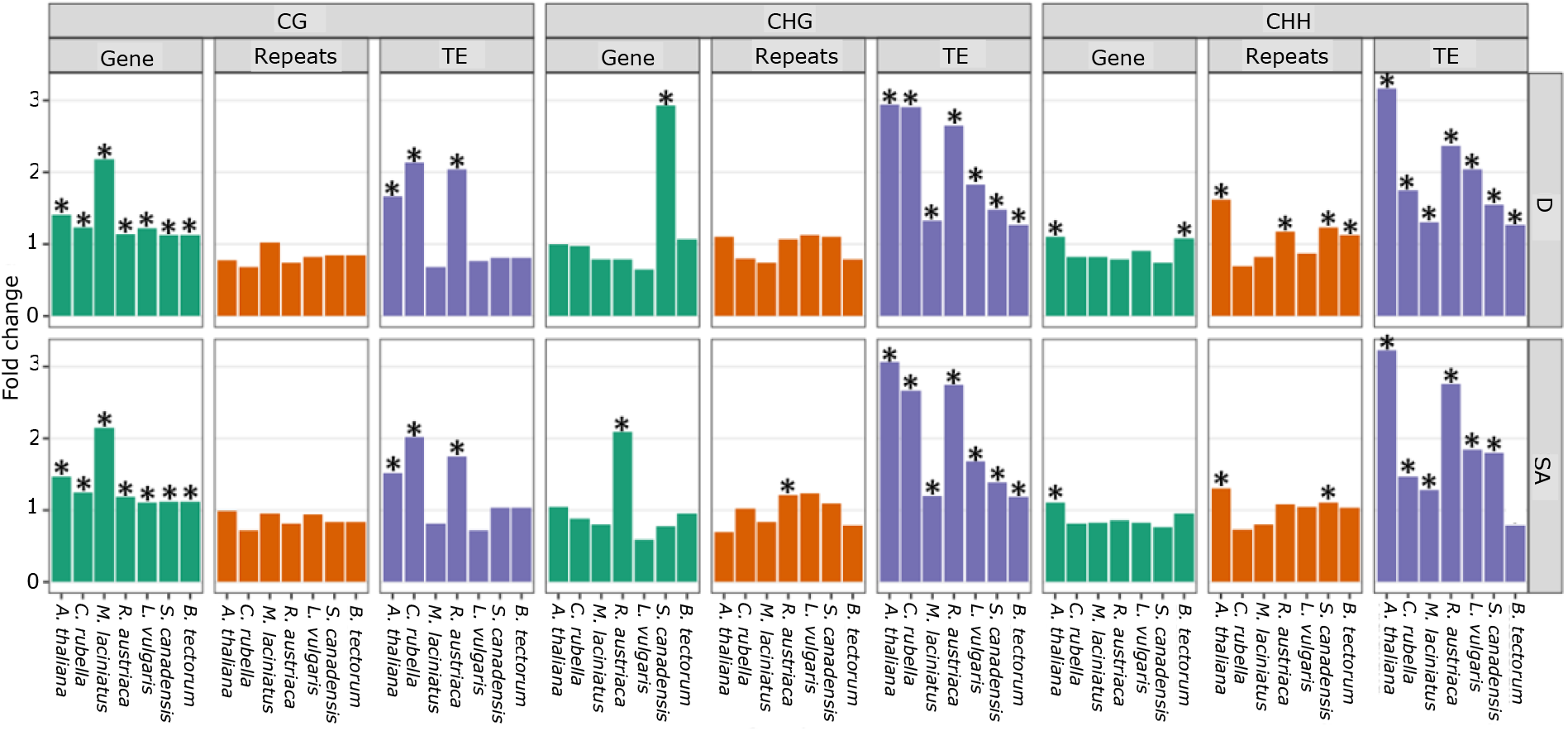
Enrichment analysis of DMCs across species, per genomic feature and cytosine context. DMCs were considered significant when the false discovery rate was < 0.05. Fold change represents the fold enrichment ratio, with a value of 1 indicating no enrichment and a value of 2 representing a 2-fold enrichment. Significance (p-value < 0.05) of fold change was evaluated using a hypogeometric test and is indicated with an asterisk (*).

## Discussion

This study evaluated methylome responses to stresses in a range of sexual and clonal plant species with different genome sizes, with the aim to establish what are similarities and differences in methylome stress-response across plant species. In all plant species, differentially methylated cytosines (DMCs) in the CG context were consistently enriched in genes, while non-CG DMCs were typically enriched in TEs. Additionally, we show that in all species all cytosines have a comparable probability to show a methylation response to stress, irrespective of sequence context. This result contrasts some recent observations that stress responses are largely biased to methylation changes in CHH context (Dubin *et al*., 2015; Wibowo *et al*., 2016). In addition to these similarities in methylome responses, clear differences in responses among species were also found. Some of these differences were associated with genome size and plant reproduction mode: large-genome species not only had more DNA methylation but also a more stress-responsive methylome (in the CG and CHH context), and clonal species showed a stronger methylation response to stress than sexual species. These specific responses may relate to the role of transposable elements in methylome plasticity and to the relative importance of plasticity in asexual lineages.

### Species differences in ‘static’ DNA methylation landscape

Confirming results from previous studies, we show that genome size is a leading factor in predicting genome-wide methylation landscape and levels (Seymour *et al*., 2014; Alonso *et al*., 2015; Niederhuth & Schmitz, 2017). This trend is attributed to the fact that differences in genome size are linked to the expansion of TEs and repeated elements; genomic features known to be heavily regulated through DNA methylation (Slotkin & Martienssen, 2007; Law & Jacobsen, 2010; Hu *et al*., 2011; Civáň *et al*., 2011). Nevertheless, genome size does not explain all variation (Vidalis *et al*., 2016). Indeed, in our study, *M. laciniatus* showed DNA methylation patterns and levels comparable to what is expected in larger genomes, despite having a relatively small genome. Similarly high methylation levels were observed in the closely related *Mimulus guttatus* (72% CG, 36.5% CHG and 6.1% CHH, genome size 321 Mb) (Colicchio *et al*., 2015), suggesting that the *Mimulus* genus might possess a relatively large number of repeats and transposable elements compared to other species. We indeed observed that a larger proportion of *M. laciniatus* epiGBS loci are annotated as TEs compared to the smaller genome-sized species (Supplementary S3). Furthermore, and similarly to what was observed in previous studies in Poaceae (Niederhuth *et al*., 2016), *B. tectorum* showed relatively low CHH methylation levels. While reduced representation techniques such as epiGBS only allow us to study small, non-targeted regions of the genome, the patterns in the methylation landscape and levels that we observed across species are in accordance with what has previously been documented (Cokus *et al*., 2008; Colicchio *et al*., 2015).

### DNA methylation stress responses

#### Methylome stress responses in different cytosine contexts

##### All cytosines contexts respond to stress

It is widely believed that cytosine methylation in different sequence contexts (CG, CHG and CHH) responds differently to environmental stimuli, as methylation of these contexts is regulated by very different enzymatic pathways (Richards, 2006; Zhang *et al*., 2018; Gallego-Bartolomé, 2020). Studies in a variety of plant species, including *Populus nigra, Thlaspi arvense* and *Arabidopsis thaliana*, suggest that CHH-type of methylations are more responsive to environmental conditions, whereas CG (and to a lesser extent CHG) are more stable and more often reflecting the genotypic origin of a plant rather than its environmental conditions (Dubin *et al*., 2015; Galanti *et al*., 2022; Peña-Ponton *et al*., 2022). Nevertheless, counter-examples exist such as in the common Duckweed *Lemna minor* or *Fragaria vesca*, where exposure to stress induced a large number of changes in methylation in the CHG and/or CG context rather than CHH (López *et al*., 2022; Van Antro *et al*., 2022). Studies that consider single-base resolution methylome responses to stress almost exclusively look at the absolute number of induced DMCs. In the present study, we took a different approach. By recording the proportion of cytosines that respond to a stress, not the absolute numbers, our results suggest that all cytosines are equally likely to show a methylation response to stress, irrespective of sequence context. Thus, the relationship between cytosine context responsiveness and stress may be more complex than what has previously been reported. Previously reported stress responses occurring primarily in CHH are likely biased by the fact that there are many more CHH-cytosines than CG or CHG-cytosines present in genomes ((López *et al*., 2022; Peña-Ponton *et al*., 2022) and as seen in our results), increasing the probability of detecting significant CHH methylation changes.

##### Enrichment of DMCs occur in genes and TEs

The methylation responses in specific cytosine contexts differ between specific genomic contexts, however. Our results clearly demonstrate that in all species, significant enrichment of CG-DMCs occurs in genes, with non-CG types of DMCs typically enriched in TEs. Such enrichment results add evidence to the growing literature that DNA methylation is associated not only with regulating gene and TE activity during normal plant functioning, but with their activity during stress responses (Mirouze & Paszkowski, 2011; Sahu *et al*., 2013; Tricker, 2015; Negi *et al*., 2016; Zhang *et al*., 2018; Kumar & Mohapatra, 2021).

Many TEs are heavily methylated in all three cytosine contexts in order to suppress TE activity and to maintain genome integrity (Slotkin & Martienssen, 2007; Civáň *et al*., 2011). Under stress conditions, we show that modification of the methylation state of TEs is a conserved response across plant species. This altered methylation state in TEs could result in their activation or repression, which in turn could allow access to stress-response regulatory sequences; modify gene expression of nearby genes and, in some cases, genetic variability might be increased via transposition (Negi *et al*., 2016; *Horváth et al*., *2017*; Roquis *et al*., 2021). Such processes might contribute to plant strategies when coping with periods of environmental stress (Mirouze & Paszkowski, 2011; Rey *et al*., 2016).

Enrichment of CG-DMCs in gene-bodies has been recorded in numerous species but the functional relevance of this response to stress in gene-bodies remains unclear (Bewick *et al*., 2019). Some studies have correlated changes in gene-body methylation to gene expression flexibility (Entrambasaguas *et al*., 2021; Wang *et al*., 2021) while others have proposed that gene-body methylation may be but a passive byproduct of other epigenetic processes in genes (Wendte *et al*., 2019). While functionality remains unclear, our study clearly demonstrates that enrichment of cytosine context specific DMCs in specific genomic regions is a highly conserved response across plant species when faced with stress.

#### Species- and Stress-specific responses

While several aspects of the methylome response to stress were shared by most, if not all, species, stress- and species-specific responses were also detected. Studies have demonstrated the effect of drought and SA on the methylomes is plant species such as *A. thaliana, Oryza sativa, Morus alba, Populus trichocarpa* and *Brassica juncea* (Dowen *et al*., 2012; Li *et al*., 2020; Liang *et al*., 2014; Sadhukhan *et al*., 2022; van Dooren *et al*., 2020; W. Wang *et al*., 2016). We further find that drought may induce a more systemic change in DNA methylation levels across the whole genome, while SA may act more at the regional or individual cytosine level. We hypothesise that our SA treatment may not have been perceived as a serious enough stress to warrant widespread and systematic changes at the molecular level, similar to the drought stress. One could expect certain species to be more resistant to certain stresses and would need a substantially higher dose to induce changes in their DNA methylation patterns (Wang *et al*., 2016).

Although the effect of stress at the genome-wide level remained limited, in all species, both drought and SA induced methylation responses at individual cytosines that were unique to that respective stress, indicating stress-specific responses at the nucleotide-specific level. But many DMCs were induced by both stresses, or in interaction between the two stresses. This would indicate that a large number of induced DMCs, irrespective of cytosine context, are sensitive to multiple environmental triggers and are thus perhaps part of more generic stress response processes rather than a stress-specific response. This is in agreement with recent observations in *P. nigra*, where different stresses often caused similar DNA methylation changes in the genome (Peña-Ponton *et al*., 2022). For the stress-specific DMCs, a relevant question is if these occur in genes (or their regulatory regions) that are functionally involved in the stress response. One limitation to using reduced-representation techniques such as epiGBS in plants is the limited opportunity to infer functional relevance from detected DMCs (Paun *et al*., 2018). Follow-up studies are needed to verify if stress-specific DMCs are found in or near stress-specific genes, while general-responding DMCs are found in genes that are involved in a general stress response. Such studies would help in elucidating the potential functional role of stress-induced DNA methylation changes.

Interestingly, *A. thaliana* often showed divergent methylome responses compared to other species. *A. thaliana* is often used as a model species for genetic and epigenetic studies, which has been invaluable for unravelling molecular mechanisms underpinning epigenetic variation.

However, in addition to results obtained previously based on multi-species comparisons of static methylomes that identified the *A. thaliana* methylome as an outlier compared to other species (Alonso *et al*., 2015; Niederhuth *et al*., 2016), our study revealed that DNA methylation stress responses of *A. thaliana* is also often different from that of other species. Specifically, we showed that a relatively low proportion of cytosines responded to stress in *A. thaliana’s* methylome. Additionally, less than 1% of *A. thaliana*’s induced DMCs showed a methylation percentage point difference of 20% or more. A possible explanation to *A. thaliana*’s lack of response could be due to its relatively low genomic TE content ((de La Chaux *et al*., 2012); Supplementary S3). Because the methylation stress response is biased to TEs, the low TE content of the *A. thaliana* genome may explain a lower overall methylation response to stress in this species.

#### Effect of Genome size and Reproduction mode

We performed our study as a series of species-level experiments that were carried out in parallel, but not as one large fully randomised experiment. This limits opportunities for direct statistical testing of species differences. However, because all species-level experiments were carried out within an 8-week period and under the same environmental conditions, and in no particular order with respect to species genome size or reproduction mode, we can interpret associations between these characteristics and methylome responses at the species level. First, species with larger genomes showed a more responsive methylome: the proportion of responding cytosines in CHH context increased with genome size. This is presumably related to a larger genomic representation of TEs in large genomes and the presence of CHH islands (Civáň *et al*., 2011; Niederhuth et al., 2016). While TEs are enriched in CHH DMCs in all species, a larger TE fraction (and thus CHH islands) in the genome can lead to a larger fraction of the genome responding to stress. Despite having a lower genomic fraction of TEs, smaller-genome species showed significant enrichment in TEs of DMCs in all three cytosine contexts, while only non-CG DMCs were enriched in larger-genome species. The reason for this is unknown, but it might reflect stronger TE suppression in smaller-genome species.

Another interesting observation is the large variation between species in the proportion of DMCs that show a strong methylation response to the stress treatment (DMCs with a methylation percentage point difference of 20 or higher). Clonally reproducing species showed significantly stronger responses of DMCs (and thus higher degree of methylome plasticity) than sexually reproducing species. Strong methylation responses to stress are more likely to have functional consequences in mediating stress responses (Anastasiadi *et al*., 2018) and are also more likely to show long-term stability in clonal lineages (Van Antro *et al*., 2022). If the strongly responding DMCs reflect functional regulation of the stress response (which our DNA methylation study does not test), then the observation is consistent with the idea that asexual lineages are selected for high phenotypic plasticity (Lynch 1984).

## Conclusion

Species-wide comparison of static methylomes have been conducted in previous studies, however, to our knowledge, this paper is the first to report a species-wide comparison of methylome stress-responses. While large discrepancies exist between studies that focus on only one species, our experiment allowed us to demonstrate that important generalities in stress responses exist among plant species. Generalities relate to bias of the stress response in TEs (particularly in non-CG contexts) and to genes (in CG context). Another generality across species that is suggested by our results is that all cytosines are equally likely to respond to stress, irrespective of cytosine context. This contrasts with recent ideas that cytosines in CHH context are more stress-responsive; we suggest that methylation in CHH cytosines only seems more responsive because there are more CHH-cytosines in the genome than CHG- or CG-cytosines. In addition to generalities, we also exposed notable differences in the methylome response to stress between species. The association of differences in methylome responses with genome size and also with plant reproductive mode can provide new insights in the potential role of DNA methylation in plant stress responses. Further insight in this role will require more functional analyses (for instance, how do observed methylation changes relate to gene expression changes?). By exposing both generalities and specifics of plant methylome responses to stress, our study shows both the strengths and the limitations of using single model species for plant DNA methylation analysis, and suggests that it is important to acknowledge ecological and life history differences in plant DNA methylation analysis.

## Acknowledgments

Cristian Pena is acknowledged for his critical questions, discussion and contributions to the experiment and data analysis. Both Cristian Pena and Paolla Rallo are thanked for their help in sampling and plant care-taking when needed. Christa Mateman and colleagues at NIOO-KNAW are acknowledged for their critical thinking and contribution during the laboratory phase of the experiment. Gregor Disveld, Eke Hengeveld and other NIOO-KNAW caretakers for maintenance and up-keep of the greenhouse facilities. The data analysis and journal submission were enabled by the European Training Network “EpiDiverse”, which received funding from the EU Horizon 2020 program under Marie Skłodowska-Curie grant agreement No 764965. All experiments, data analysis and writing was conducted at NIOO-KNAW. This publication is filed under NIOO-KNAW publication number XXX

## Author Contributions

MVA, KJFV, PV and LM designed the experiment. SI and MVA propagated the plants, collected samples and conducted the laboratory aspect of the experiment. MP provided support with the epiGBS pipeline and data analysis. MVA, KJFV, PV, LM and WHP performed the statistical analysis presented in this paper. The manuscript was drafted by MVA, KJFV, PV and WHP, with input and approval of the final version by all authors.

## Data Availability

Please email authors if you would like access to the data. The data will become accessible through Zenodo, NCBI and github once the manuscript has been peer-reviewed.

