## Supplementary for "DNA methylation responses to stress across different plant species"

### Supporting Information

***Supplementary data S1: Accession and origin of seeds used to propagate G1 parents***

| Species | Accession Number | Origin |
|---|---|---|
| A. thaliana | Col 0 - MA line, 31-29-6 | Provided by Claude Bekker, LMU BioCenter |
| C. rubella | Cr1GR1 | Provided by Tanja Slotte, Stockholms University |
| M. laciniatus | SNB CFT-3 | Provided by Jack Colicchio, UC Berkley |
| F. vesca | FV_CR_05_11_S1_180801_C-1 | Provided by Iris Sammaco, EpiDiverse |
| R. austriaca | - | NIOO-KNAW |
| L. vulgaris | 595510 | Vreeken's Zaden |
| S. canadensis | 345195/604720 | Vreeken's Zaden |
| B. tectorum | PI 211003 | GRIN US Germplasm W6 |

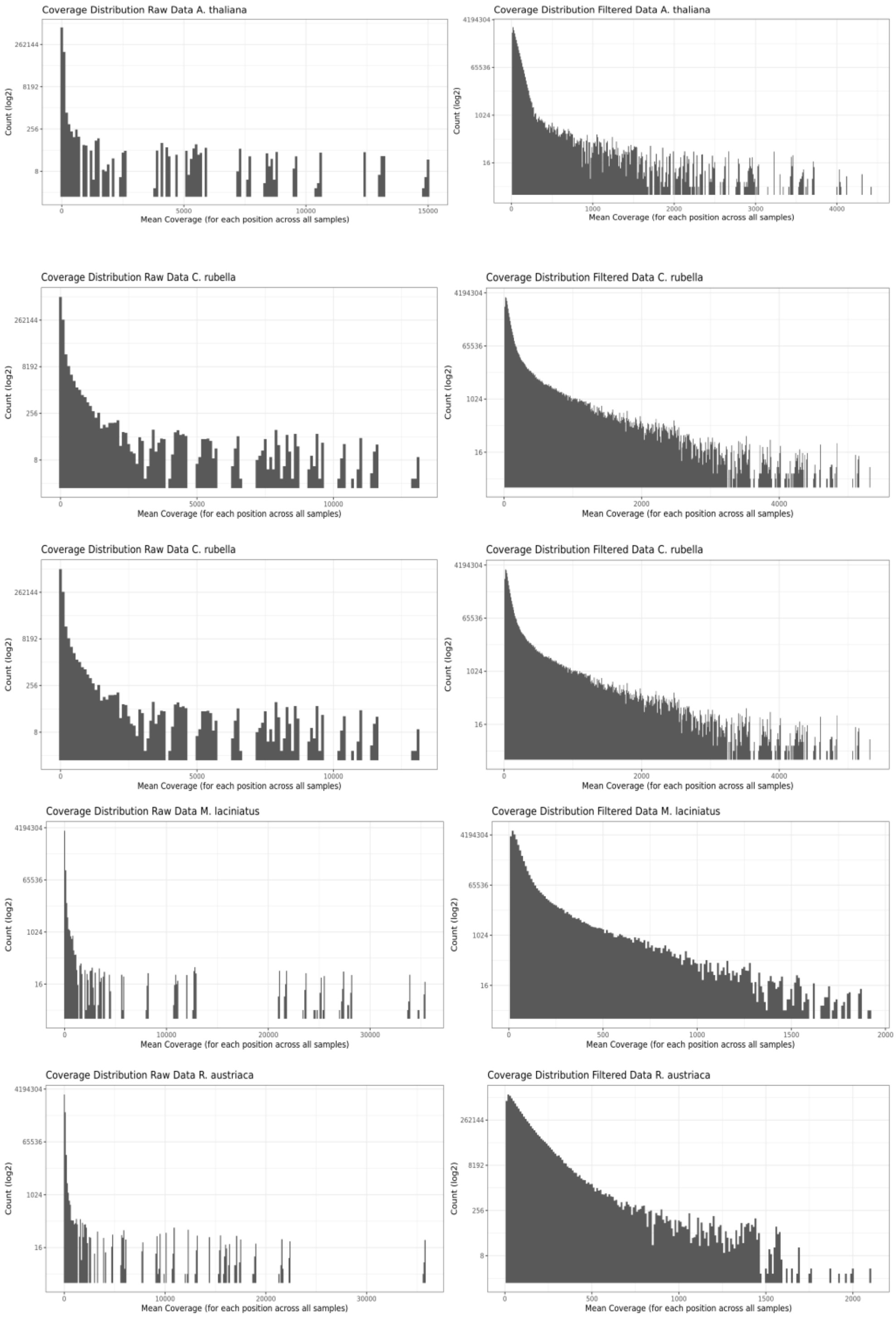

Coverage Distribution Raw Data A. thaliana
Count (log2)
Mean Coverage (for each position across all samples)
Coverage Distribution Filtered Data A. thaliana
Count (log2)
Mean Coverage (for each position across all samples)
Coverage Distribution Raw Data C. rubella
Count (log2)
Mean Coverage (for each position across all samples)
Coverage Distribution Filtered Data C. rubella
Count (log2)
Mean Coverage (for each position across all samples)
Coverage Distribution Raw Data C. rubella
Count (log2)
Mean Coverage (for each position across all samples)
Coverage Distribution Filtered Data C. rubella
Count (log2)
Mean Coverage (for each position across all samples)
Coverage Distribution Raw Data M. laciniatus
Count (log2)
Mean Coverage (for each position across all samples)
Coverage Distribution Filtered Data M. laciniatus
Count (log2)
Mean Coverage (for each position across all samples)
Coverage Distribution Raw Data R. austriaca
Count (log2)
Mean Coverage (for each position across all samples)
Coverage Distribution Filtered Data R. austriaca
Count (log2)
Mean Coverage (for each position across all samples)

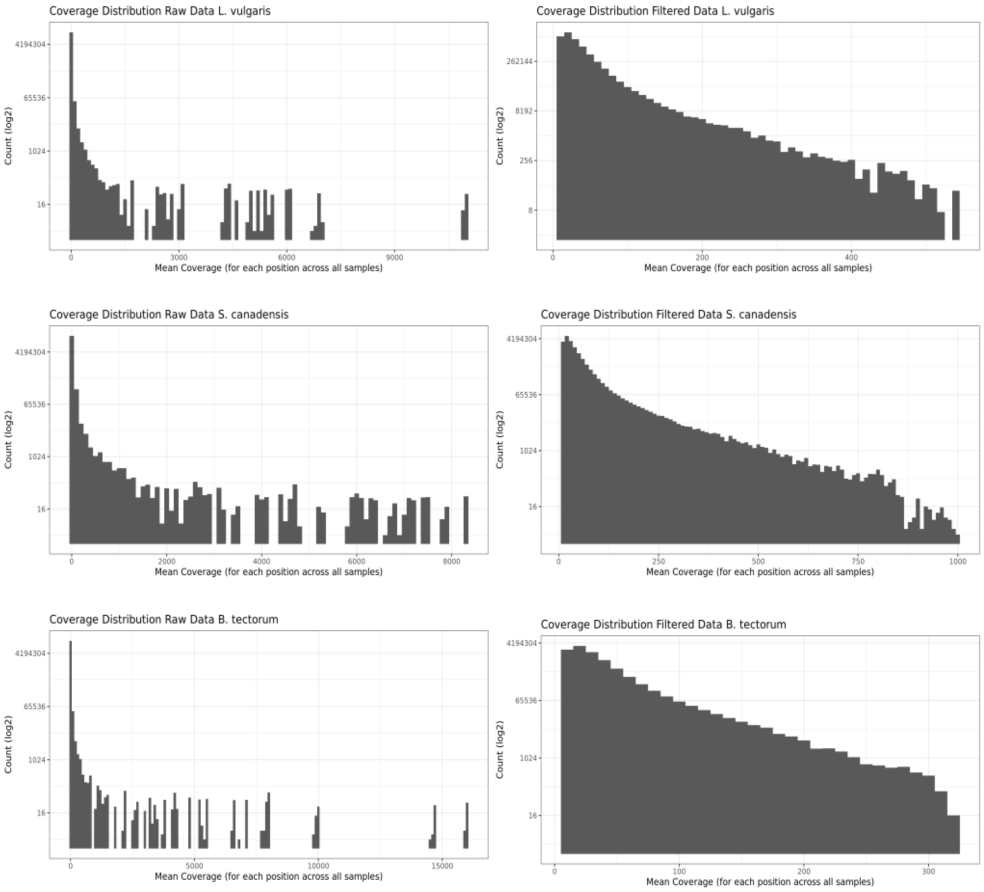


***Supplementary data S2: Mean coverage of each cytosines (across all samples)*** *of the raw sequencing data and filtered data. Filtering step included removing cytosines whose mean coverage was < 10X as well as removing the 0.1% of cytosines with the highest coverage.*

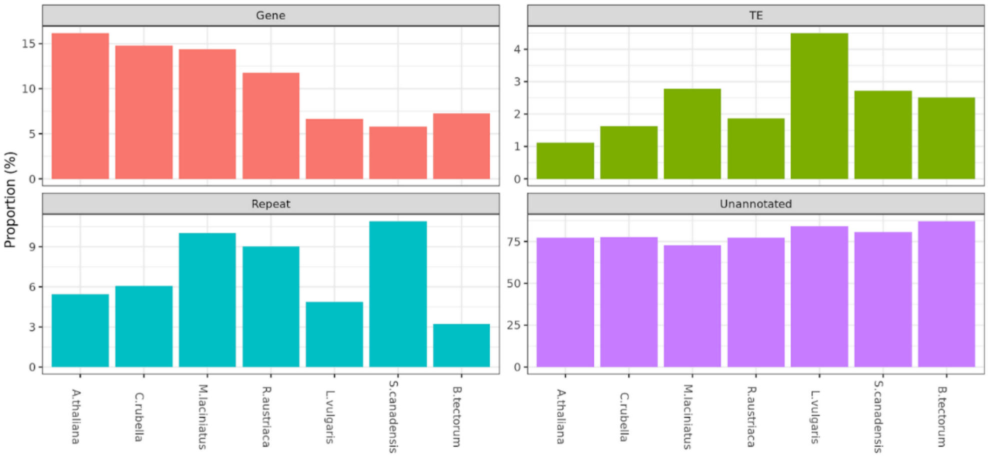


***Supplementary data S3: For each species, proportion of epiGBS loci that were annotated*** *either as Gene, Transposable element (TE), Repeats and Unannotated based on homology.*

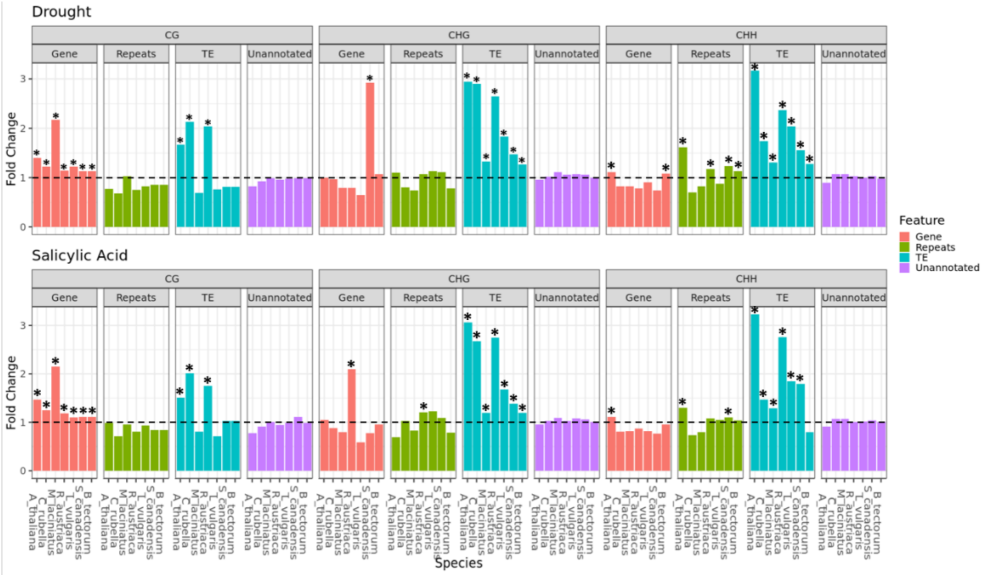


***Supplementary data S4: Enrichment analysis of DMCs across species, per genomic feature and cytosine context.*** *Including unannotated regions. DMCs were considered significant when the false discovery rate was < 0.05. Fold change represents the fold enrichment ratio, with a value of 1 indicating no enrichment and a value of 2 representing a 2-fold enrichment. Significance (p-value < 0.05) of fold change was evaluated using a hypogeometric test and is indicated with an asterisk (*).*
